# Development of an *in vitro* pre-mRNA splicing assay using plant nuclear extract

**DOI:** 10.1101/222976

**Authors:** Mohammed Albaqami, Anireddy S. N. Reddy

## Abstract

**Background:** Pre-mRNA splicing is an essential post-transcriptional process in all eukaryotes. *In vitro* splicing systems using nuclear or cytoplasmic extracts from mammalian cells, yeast, and *Drosophila* have provided a wealth of mechanistic insights into assembly and composition of the spliceosome, splicing regulatory proteins and mechanisms of pre-mRNA splicing in non-plant systems. The lack of an *in vitro* splicing system prepared from plant cells has been a major limitation in splicing research in plants.

**Results:** Here we report an *in vitro* splicing assay system using plant nuclear extract. Several lines of evidence indicate that nuclear extract (NE) derived from *Arabidopsis* seedlings can convert pre-mRNA substrate (*LHCB3*) into a spliced product. These include: i) generation of an RNA product that corresponds to the size of expected mRNA, ii) a junction-mapping assay using S1 nuclease revealed that the two exons are spliced together, iii) the reaction conditions are similar to those found with non-plant extracts and iv) finally mutations in conserved donor and acceptor sites abolished the production of the spliced product.

**Conclusions:** This first report on the plant *in vitro* splicing assay opens new avenues to investigate plant spliceosome assembly and composition, and splicing regulatory mechanisms specific to plants.

## Background

Shortly after the discovery of introns in 1977, different groups developed the mammalian cell-free system (*in vitro*), using nuclear or cytoplasmic extracts, which were competent for pre-mRNA splicing [1-3]. Subsequently, the preparation of efficient splicing extracts and the *in vitro* splicing assay have been adapted to other organisms such as budding yeast (*Saccharomyces cerevisiae*) and fruit flies (*Drosophila melanogaster*) [4, 5]. The development of mammalian, yeast, and *Drosophila in vitro* systems to study pre-mRNA splicing has provided essential insights into spliceosome assembly, its composition and splicing mechanisms in non-plant systems. The characterization of the splicing two-step trans-esterification reaction, pre-mRNA splicing intermediates, and the formation of final mature mRNA and lariat structure of intron have all been revealed by *in vitro* splicing studies [3, 6]. Furthermore, *in vitro* splicing combined with immunodepletion has long been used to determine the roles of spliceosomal components, such as small nuclear ribonucleic proteins (snRNP) [7, 8]. Several other splicing regulators were characterized by their ability to promote the *in vitro* splicing assay [9, 10]. Furthermore, the formation of spliceosomal complexes and their stepwise assembly pathway have been deduced *in vitro* by native (nondenaturing) agarose/polyacrylamyde gel electrophoresis [11-13]. Additionally, many *in vitro* biochemical splicing studies have allowed purification of spliceosomes and provided a wealth of information on the spliceosome’s composition, its structures, and the structure conformational dynamics of spliceosomal complexes [13-21]. The identification of additional splicing regulatory *cis*-elements, such as splicing enhancers or silencers and their cognate factors, has also been expedited by studies using *in vitro* biochemical assays [22-27]. The *in vitro* splicing assay has also been a valuable technique for dissecting abnormal splicing events, which cause human genetic disease, and in developing new therapeutic approaches for human disease [2, 28]. Finally, the *in vitro* splicing assay has been used to evaluate molecules with splicing inhibitory functions, such as spliceostatin A [29]. These are some examples to illustrate how *in vitro* splicing assays have contributed to important information on regulatory mechanism of gene expression. The *in vitro* splicing systems have been, and will continue to be, indispensable tools for studying the mechanism of splicing.

As in animals, pre-mRNAs from a majority (over 80%) of plant genes contain non-coding sequences and are processed to generate mature mRNAs [30, 31]. Recent studies indicate that the developmental and environmental cues can reprogram gene expression in plants by regulating post-transcriptional processes, especially pre-mRNA splicing [31-38]. However, many aspects of pre-mRNA splicing in plants are yet to be elucidated. Furthermore, the composition of the plant spliceosome and its assembly intermediates are currently undefined [39, 40]. Thus, the study of pre-mRNA splicing in plants requires innovative approaches, which will greatly empower this field of research.

In spite of the absence of an *in vitro* plant-splicing system, efforts have been made to identify plant spliceosomal components. Bioinformatics analyses using sequence similarity have identified the core components of the plant spliceosome, including five snRNAs and several orthologs of known spliceosomal proteins [40-44]. Likewise, the highly conserved sequences at the 5′ splice site (5′ss), 3′ splice site (3′ss), polypyrimidine tract and branch point sequence (BPS) are similar between plants and animals [31]. These similarities and some reports on splicing of plant introns in mammalian splicing extracts [45, 46] suggest similar basic mechanisms of intron processing across eukaryotic systems, but there are also numerous indications of plant-specific splicing regulatory mechanisms. For example, animal introns cannot be processed or animal transcripts are aberrantly spliced in plant systems [47, 48], the average sizes of plant introns are shorter than their mammalian counterparts [40], and analysis of proteins similar to mammalian spliceosomal proteins indicates that there is almost twice the number of plant splicing factors compared to human splicing factors [40, 44]. Other studies have also shown that plants and animals use different mechanisms to recognize splice sites, especially 3’ splice sites [49-52]. Furthermore, comparative analysis of alternative splicing (AS) events between plants and animals has revealed that intron retention is the predominant mode of AS in plants [53, 54], whereas exon skipping is the predominant mode in animals [55]. Although the pre-mRNA splicing mechanisms in plants are poorly understood, based on the reasons cited above it is likely that the mechanisms of recognition of introns and exons involve both similar and plant-specific mechanisms.

The development of *in vitro* systems to study RNA-related mechanisms in plants is limited and challenging (Sugiura, 1997; Reddy *et al.,* 2013). An *in vitro* pre-mRNA splicing system to uncover plant splicing regulatory mechanisms has long been awaited [39, 40]. Therefore, despite difficulties inherent to plant cells, here we describe our efforts to develop an *in vitro* system for plant pre-mRNA splicing using nuclear extract (NE) prepared from *Arabidopsis* seedlings. We present a detailed procedure for the preparation of the NE and a subsequent *in vitro* splicing reaction using a plant pre-mRNA substrate containing a single intron from *LIGHT-HARVESTING CHLOROPHYLL B-BINDING PROTEIN 3* (*LHCB3*). We show that plant NE is capable of converting *LHCB3* pre-mRNA substrate to the size of expected mRNA. This is the first step toward establishing a plant-derived *in vitro* pre-mRNA splicing assay. This study opens new avenues to investigate spliceosome assembly and composition, splicing regulatory mechanisms specific to plants, and thereby enhances the overall understanding of post-transcriptional gene regulatory mechanisms in eukaryotes.

## Materials and Methods

### *Arabidopsis* nuclear extract preparation

The nuclear extract preparation method presented here is a modification of protocols described previously [56-58]. For plant material preparation, seeds (50 mg) of *Arabidopsis thaliana* ecotype Columbia-0 (Col-0) were surface-sterilized with 70% ethanol followed by 15% bleach and stratified for 3 days at 4 °C. Then, seeds were placed into 100 mL of Murashige and Skoog (MS) medium (1x MS basal salts, 1 mL/L MS vitamin solution, and 1% sucrose, pH 5.7) in a 250 mL flask and moved to a growth chamber. Seedlings were grown in a flask on a shaker at 150 rpm in dark at 24°C for 4 days. Four-day-old seedlings were harvested, rinsed three times with Nanopure water, and excessive water was removed using a few layers of Kimwipes. Afterwards, the seedlings were weighed, directly frozen in liquid N_2_, and stored at −80°C.

For nuclear protein preparation, 5 g of etiolated seedlings were ground into a fine powder in liquid nitrogen. Subsequently, the sample was homogenized in 25 mL of Honda buffer (1.25% Ficoll 400, 2.5% Dextran T40, 0.44M sucrose, 10mM MgCl_2_, 0.5% Triton X-100, 20mM HEPES KOH, pH 7.4, 5 mM DTT, 1 mM PMSF, and 1% protease inhibitor cocktail [Sigma-Aldrich, St. Louis, MO; catalog number: P9599]) for 15 min on ice, with gentle mixing every minute. The homogenate was filtered through two layers of Miracloth into a 50 mL Corex tube. Then the residue left behind on the Miracloth was washed with 25ml ice-cold Honda buffer, and then the filtration step was repeated and collected into the same Corex tube. The filtrate (total 50 ml) was centrifuged at 2000 *g* for 15 min at 4°C. The supernatant was discarded and the pellet resuspended in 15 mL ice-cold Honda buffer, then incubated on ice for 15 min with gentle mixing every minute. It is not recommended to pipet up and down when mixing, as this can disrupt the nuclei; instead, a camel-hair brush should be used to resuspend the nuclei pellet. After complete resuspension, the sample was centrifuged at 1500 *g* for 15 min at 4°C. This washing step was repeated two times. The pellet was then resuspended in 15 mL ice-cold washing buffer (20 mM HEPES KOH, pH 7.9, 100 mM KCl, 0.2 mM EDTA, 10% (v/v) glycerol, 1mM DTT, 1 mM PMSF, and 1% protease inhibitor cocktail [Sigma-Aldrich, St. Louis, MO; catalog number: P9599]) for 15 min on ice, mixed gently every minute. It is also not recommended to mix by pipetting here. After complete resuspension, the sample was centrifuged at 1500 *g* for 15 min at 4°C. The nuclei pellet was then resuspended in 0.5X ice-cold nuclei swelling buffer (50 mM Tris–HCl (pH 7.9), 10 mM 2-mercaptoethanol, 20% glycerol, 5 mM MgCl_2_, 0.44 M sucrose, 1 mM PMSF, and 1% protease inhibitor cocktail [Sigma-Aldrich, St. Louis, MO; catalog number: P9599]) and transferred to a 1.5 ml microcentrifuge tube, then incubated at 4°C for 30 min with gentle rocking. The extract was then centrifuged for 30 min at maximum speed (16,000 *g*) at 4°C and the supernatant removed to a new 1.5 mL microcentrifuge tube. Then, the NE was distributed into 50 μl aliquots, flash-frozen in liquid nitrogen and stored at −80°C for *in vitro* splicing assay.

### DNA templates and *in vitro* pre-mRNA synthesis

For *in vitro* pre-mRNA synthesis, DNA templates were amplified by PCR from *Arabidopsis* genomic DNA, using a gene-specific forward primer plus SP6 promoter sequence and reverse primer plus an adaptor sequence. PCR products of the correct size were then gel-purified using the Thermo Scientific GeneJET Gel Extraction Kit (Thermo Fisher Scientific, Waltham, MA; catalog number: K0691). Purified DNA templates were quantified using a NanoDrop 1000, and approximately 0.250 μg of amplified DNA template was used for *in vitro* transcription. To generate mutations at conserved splicing sites, DNA template sequences with desired sequences were synthesized at Integrated DNA Technologies, Inc. (Coralville, IA; https://www.idtdna.com).

[^32^P]-labelled pre-mRNA substrates were prepared using an *in vitro* transcription system as described previously in [59]. The pre-mRNA substrates were internally labeled with 45 μCi of [α-^32^P] UTP (800 Ci mmol^−1^, PerkinElmer, Waltham, MA) using SP6 RNA polymerase (Fermentas, Thermo Fisher Scientific, Waltham, MA; www.fermentas.com) in the presence of 500 μM ATP and CTP, 50 μM GTP and UTP, 50 μM cap analog (^7m^GpppG), and 20 U RNase inhibitor. *In vitro* synthesized [^32^P]-labelled pre-mRNAs with the correct size were gel-purified using TNS solution (25 mM Tris-HCl (pH 7.5), 400 mM NaCl, 0.1% SDS) for overnight at room temperature. Radioactive pre-mRNAs were measured using a liquid scintillation counter (Tri-Carb Liquid Scintillation Counter, PerkinElmer, Waltham, MA); 25,000 CPM (~20 fmol) of [^32^P]-labelled pre-mRNA substrates was used for *in vitro* splicing reactions, unless otherwise specified (see figure legends).

### *In vitro* splicing

Unless otherwise specified, *in vitro* splicing reactions (25 μl) contained 1 mM ATP, 20 mM creatine phosphate (CP), 10 U RNase inhibitor, 1 mM DTT, 72.5 mM KOAc, 25,000 CPM (~20 fmol) [^32^P]-labelled pre-mRNA, and 50% NE. Reactions were incubated at 30°C for the times indicated in figure legends. Subsequently, 175 μl of proteinase K master mix (1× proteinase K buffer, 0.25 mg/ml glycogen, 0.25 mg/ml proteinase K, and sterile water) was added, and the solution incubated at 37°C for 20 min. Afterward, RNA was purified by adding an equal volume of phenol:chloroform, precipitating with 2.5 volumes of 100% ice-cold ethanol, and air-drying the isolated pellet for 5 min [60]. Finally, RNA samples were dissolved with formamide/EDTA stop dye (formamide with 0.1% bromophenol blue, 0.1% xylene cyanol, and 2 mM EDTA).

### Visualization of splicing products

Purified RNA from *in vitro* splicing reactions was analyzed by fractionation on a 6% polyacrylamide-urea gel as described previously [60]. RNA samples were heated at 95°C for 3 min, and loaded onto a pre-run gel. The gel was run at 200 V for 3 hours. Subsequently, the gel was transferred onto Whatman paper and dried for 2 hours using a Bio-Rad Gel Dryer at 80°C with suction. The gel was then exposed to a phosphor-imaging screen overnight, and imaged using a STORM 840 imager (Molecular Dynamics, GE Healthcare, Little Chalfont, United Kingdom; www.gehealthcare.com).

### S1 nuclease protection assay

For the S1 nuclease protection assay, *in vitro* spliced RNA (spliced RNA) was gel-purified as described above. The spliced product from at least three samples was pooled together. DNA oligo (50 nt, 100 μM) with sequence complementary to the exon/exon junction was hybridized to the purified RNA in S1 hybridization buffer (80% formamide, 40mM PIPES (pH 6.4), 500 mM NaCl, 1 mM EDTA) for 2 hours at room temperature after denaturing at 95°C for 5 min [61].Subsequently, total hybridized [^32^P]-labelled RNA/oligo DNA was digested with S1 nuclease (100U) (Promega, Madison, WI; catalog number: M5761) in 1x S1 nuclease buffer (provided with the enzyme) for 1 hour at 37°C. The undigested [^32^P]-labelled RNA was purified by phenol:chloroform extraction and visualized as described above.

## Results

### *Arabidopsis* NE processed *LHCB3* pre-mRNA to produce an expected size mRNA

Generally, *in vitro* splicing reactions are conducted using NE prepared from HeLa cells; however, there are reports of NEs from other cells, such as *Drosophila* Kc cells, also being used [62]. The quality of NE is a vital factor for a successful and efficient in *vitro* splicing assay [62]. Therefore, we aimed first to develop a method for the preparation of NE from *Arabidopsis* etiolated seedlings to use in *in vitro* splicing assays. We tried three different modified methods of NE preparation from either etiolated or light-grown seedling. Only one of these methods that is modified from different protocols [56-58] using etiolated seedlings was found to be competent in splicing a pre-mRNA. This NE preparation protocol is described in detail in the methods section (Figure S1). The other two modified methods that use i) hexylene glycol-based buffers followed by Percoll density gradients [56] or ii) Honda buffer method[63] either did not show splicing of pre-mRNA or showed very weak splicing activity in our hands. Hence, these methods are not described in detail.

To test this *Arabidopsis* NE for *in vitro* splicing activity, we used a pre-mRNA substrate that contained a single intron flanked with two truncated exons from *Arabidopsis* gene light harvesting complex B3 (*LHCB3*) (Figure 1A and B and S2). The *LHCB3* pre-mRNA substrate carried an 82 nt 5′ exon, 64 nt intron, and 51 nt of 3′ exon plus a 22 nt adaptor sequence for a total length of 219 nt (Figure 1C). The [^32^P]-labelled *LHCB3* pre-mRNA was incubated at 30°C in *Arabidopsis* NE for 0, 90, or 180 min. The reaction products were isolated and analyzed by 6% denaturing polyacrylamide gel (Figure 2A). We did not observe any [^32^P]-labelled spliced RNA at 0 min incubation time; however, at 90 and 180 min there were multiple [^32^P]-labelled species smaller than the pre-mRNA substrate. The predicted spliced mRNA is 155 nt long (Figure 1C). Certainly, a 155 nt RNA, the expected size of spliced mRNA, was observed after 90 min, and this product was accumulated to a higher level after 180 min (Figure 2A). We also generated a [^32^P]-labelled marker mRNA from an *LHCB3* cDNA template (M^*^) for comparison with the *in vitro* product. In addition, we observed other RNA species (indicated with asterisks) consistently, which are likely to correspond to the pre-mRNA splicing intermediates or degradation products (Figure 2A). This finding suggests the possible *in vitro* splicing of *LHCB3* pre-mRNA using *Arabidopsis* NE.

**Figure 1:**
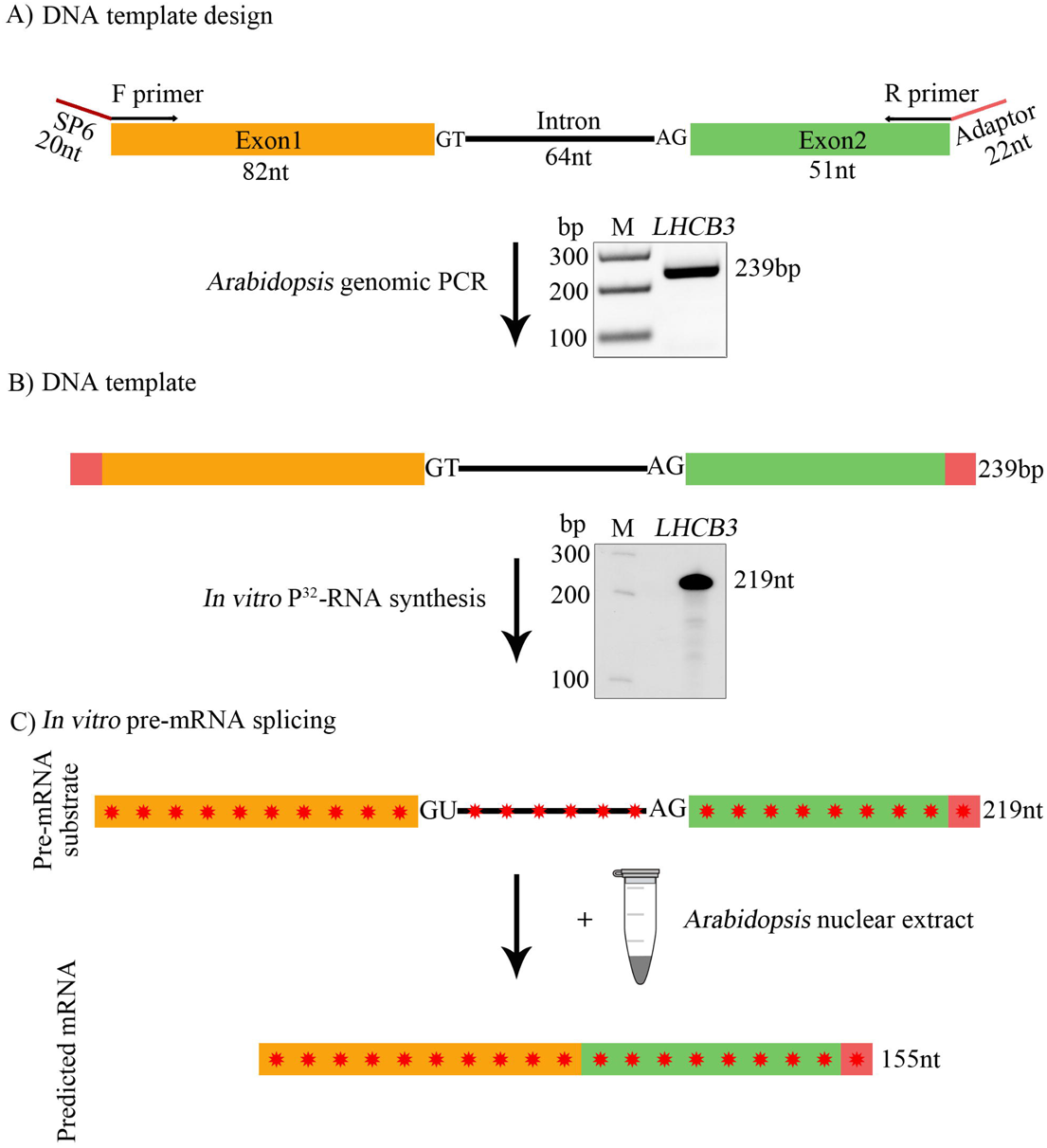
Preparation of *LHCB3* [^32^P]-labeled pre-mRNA substrate used for *in vitro* splicing assay. (A) Top, A schematic representation of a region of *Arabidopsis LHCB3* (AT5G54270) gene used to prepare DNA template to synthesize pre-mRNA substrate. A portion of the third and fourth exons (Orange and green boxes, respectively, labeled as exon 1 and exon2) and second intron (black line, labeled as intron) was used. F primer, forward primer with SP6 promoter sequence (red line), R primer reverse primer with an adaptor sequence (red line). Bottom, PCR fragment amplified with F and R primers using *Arabidopsis* genomic DNA. The PCR product was gel purified and used for *in vitro* transcription reaction. (B) Top, schematic of DNA template that was used to synthesize [^32^P]-labeled *LHCB3* pre-mRNA substrate. Bottom, A representative autoradiogram of *in vitro* [^32^P]-labeled *LHCB3* pre-mRNA substrate. (C) Description of *in vitro* splicing assay. Top, Schematic representation of labeled *LHCB3* pre-mRNA substrate used for *in vitro* splicing assay. Bottom, predicted mRNA after *in vitro* splicing of pre-mRNA substrate. Sizes of intron, exons, pre-mRNA, and predicted mRNA are indicated. Red asterisks indicate [^32^P]-nucleotides in RNA.

**Figure 2:**
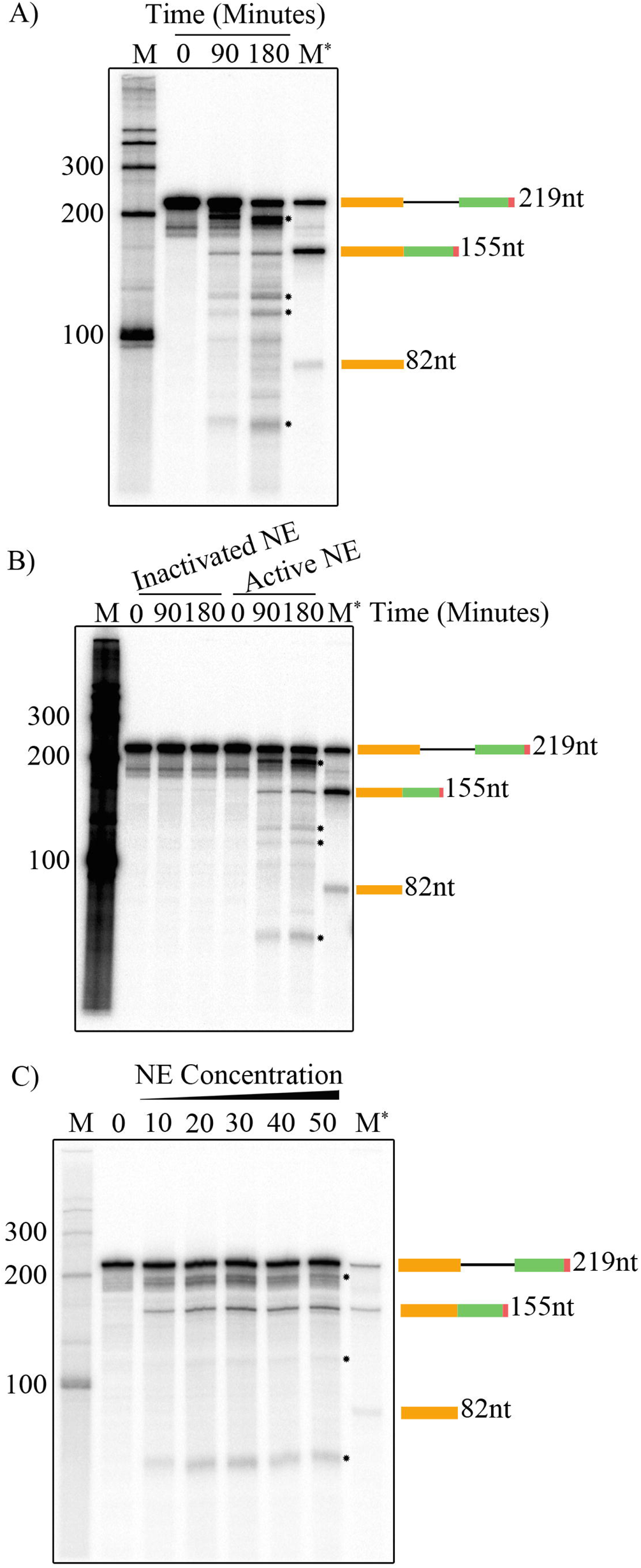
*In vitro* splicing assays. (A) *In vitro* splicing assay with the *Arabidopsis LHCB3* pre-mRNA substrate. Radioactive *LHCB3* pre-mRNA substrate was synthesized *in vitro* with a DNA template using SP6 RNA polymerase (see Fig. 1B) as described in materials and methods. [^32^P]-labeled *LHCB3* pre-mRNA substrate (25,000 cpm) was incubated with nuclear extract from *Arabidopsis* etiolated seedlings at 30°C as described in materials and methods. Samples were withdrawn at intervals (0, 90 and 180 min), [^32^P]-RNA was extracted and analyzed by electrophoresis on a 6% polyacrylamide gel containing 7 M urea, The gel was dried and exposed to a phosphor-imaging screen. (B) Heat-inactivation of *Arabidopsis* NE abolished the production of a spliced product. NE from *Arabidopsis* etiolated seedlings was incubated at 90°C for 3 min or kept on ice (as a control) were used for splicing assays at 30°C with the *LHCB3* [^32^P]-pre-mRNA. Samples were withdrawn at different time points (0, 90 and 180 minutes), [^32^P]-RNA was extracted and analyzed as described above. (C) Spliced product is increased with increasing nuclear extract concentration. In vitro splicing of [^32^P]-labeled LHCB3 pre-mRNA substrate (25,000 cpm) was carried out at 30°C in 25 μl reaction volume containing different concentrations 0-50% (v/v) of nuclear extract as described in materials and methods. All reactions were stopped after three hours; [^32^P]-RNA was extracted and analyzed as described above. M indicates [^32^P]-labeled RNA markers synthesized *in vitro* using RNA Century™-Plus Marker Templates (Applied Biosystems, AM7782). M lane contains [^32^P]-labeled *LHCB3* pre-mRNA, spliced mRNA, and exon1. Schematic diagrams on the right show pre-mRNA, spliced mRNA and exon 1 and their sizes. One of the [^32^P]-RNA products formed in *in vitro* splicing assay corresponds to the size of spliced [^32^P]-mRNA marker, suggesting that it could be a spliced product. The asterisks indicate the potential splicing intermediates. Other [^32^P]-RNA products could be another pre-mRNA splicing intermediates and/or degradation products.

### Heating of *Arabidopsis* NE inactivated *in vitro* splicing reaction

Pre-mRNA splicing is mediated via the spliceosome, which is a large and dynamic machine containing five snRNPs and numerous proteins [13]. It has been reported that heat-treated NE prepared from HeLa cells or yeast whole cell extract was unable to form a functional spliceosome and splice pre-mRNA *in vitro* [64, 65]. Therefore, we incubated the *Arabidopsis* NE at 90°C for 3 min, and then tested its splicing activity with [^32^P]-labelled *LHCB3* pre-mRNA substrate. While the untreated NE converted the input pre-mRNA to the expected product, heat-treated NE was unable to do so (Figure 2B). In agreement with non-plant splicing extracts, these results suggest that the *Arabidopsis* NE contains heat-sensitive components that are required for producing the splicing product.

### Spliced product increased with increasing NE concentration

In order to determine the effect of NE concentration on the production of the spliced mRNA, a range of NE concentrations from 0 to 50% (v/v) were tested while maintaining consistent volume and chemical composition of the reactions by adding an appropriate amount of NE-containing buffer. Indeed, the production of the splicing product increased with increasing NE concentration (Figure 2C). In addition, it seems that 30% (v/v) is an optimal NE concentration for this *in vitro* splicing assay. Thus, these results support that the appearance of the spliced mRNA is dependent on the concentration of proteins present in the NE.

### Characterization of spliced product using S1 nuclease protection assay

To determine whether the [^32^P]-labelled *LHCB3* pre-mRNA substrate was accurately spliced *in vitro,* the spliced product was analyzed by S1 nuclease protection assay. For this assay, we used a DNA oligonucleotide (50nt) probe designed to pair with the exon junction that is predicted to be joined together during *in vitro* splicing (Figure 3A). The [^32^P]-labelled spliced product generated in the *in vtiro* assay was gel-purified and at least three samples pooled together to enhance the amount of RNA present. Probes were allowed to hybridize with the purified RNA sample. In addition, the probe was also hybridized with the unspliced [^32^P]-labelled *LHCB3* pre-mRNA substrate as a negative control. Subsequently, the hybridized molecules were digested with single-strand nucleic acid-specific S1 nuclease. The S1-resistant products were then separated on a denaturing polyacrylamide gel and visualized by autoradiography (Figure 3B). The estimated size of the protected sequence is ~50 nt long, and indeed, the predicted *in vitro* splice product generated S1-resistant product with approximately size of ~50 nt and smaller products of sizes of around 25nt. Conversely, the unspliced pre-mRNA generated only S1-resistant products with a size of 25 nt that correspond to unligated exon regions on each side of the intron. These results suggest that the *LHCB3* pre-mRNA is accurately spliced *in vitro* to generate the authentic mRNA sequence spanning the *LHCB3* exon1-exon2 junction.

**Figure 3:**
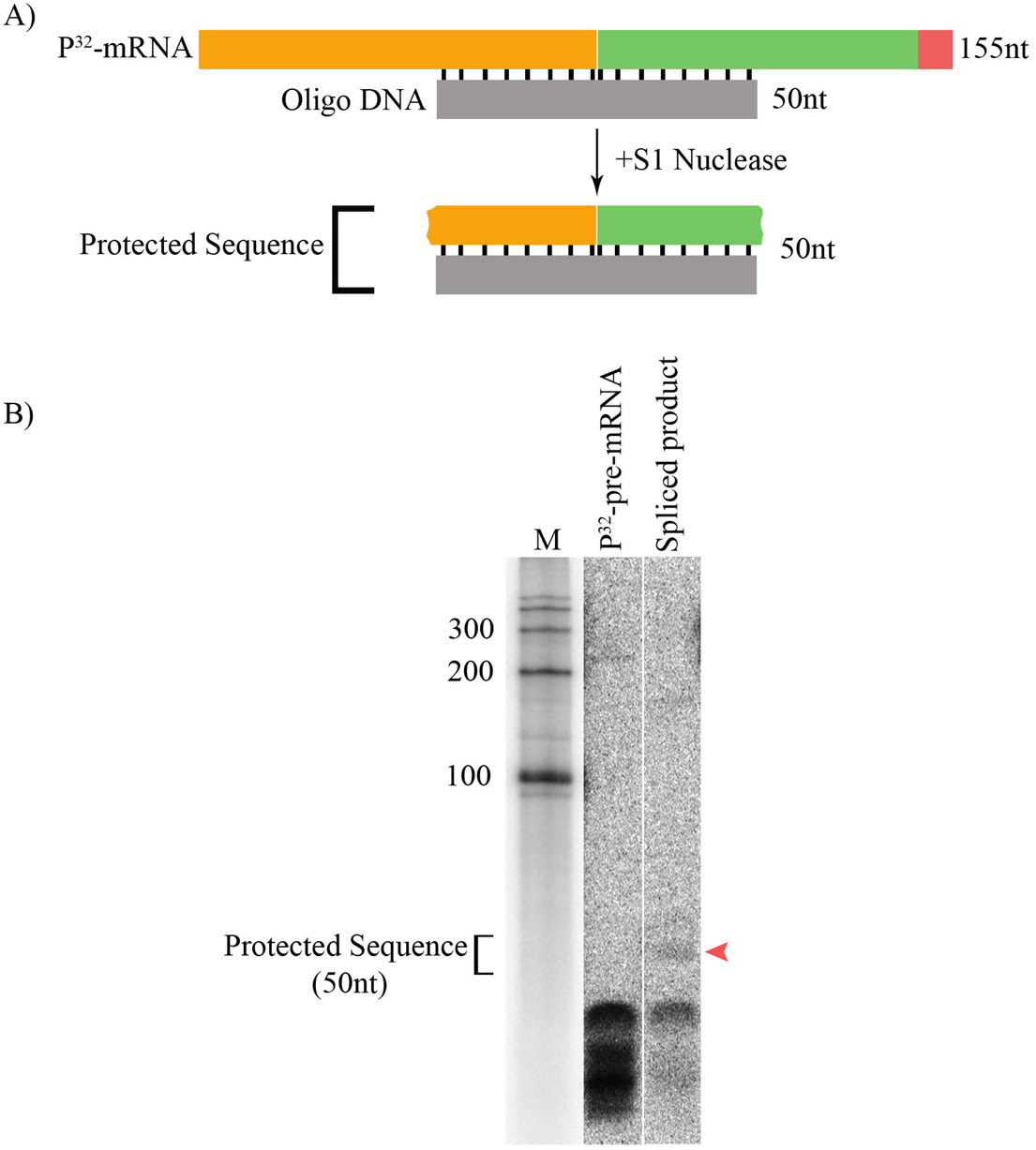
Characterization of the spliced product using S1 nuclease. (A) Schematic representation of S1 nuclease protection assay. Top, diagram of hybrid formed between spliced RNA and DNA oligonucleotide (50nt) complementary to exons junction. Bottom, diagram of protected sequences after S1 nuclease digestion. (B) Spliced [^32^P]-RNA produced in *in vitro* splicing assay was gel purified as described in material and methods. [^32^P]-pre-mRNA (negative control) and spliced product [^32^P]-RNA were hybridized to oligo DNA that is complementary to exons junction. Following hybridization, [^32^P]-RNA-DNA hybrids were digested with S1 nuclease that degraded single stranded nucleic acids. The size of the protected region (50nts) is indicated. Red arrowhead shows protected exon junction sequence with spliced product.

### Mutations in conserved splice sites modulate the production of spliced product

During the pre-mRNA splicing process, the spliceosome assembles around the 5′ss at the beginning of an intron and the 3′ss at the end of that intron [14]. It is well-established that each splice site in plants and animals consists of consensus sequences, and these include almost invariant dinucleotides: GT at the 5′ss and AG at the 3′ss [31]. *In vitro* mutation analysis of the conserved 5′ss GT revealed that these mutations could modulate splice site choices and reduce the rate of two exons ligating [66]. Therefore, we aimed here to investigate if substitution mutations of 5′ss GT in the *LHCB3* pre-mRNA substrate affect the production of its *in vitro* spliced product. To this end, we generated a *LHCB3* pre-mRNA substrates carrying substitution mutation of 5′ss GT to AC (Mutant 1, M1) (Figure 4A). In addition, since a previous study showed that mutation of the 5′ss GU did not prevent cleavage in 5′ss region, but only affected the production of spliced mRNA {Aebi, 1987 #18615}, we also included another mutated pre-mRNA substrate (Mutant 2, M2). In this substrate, both 5′ss and 3′ss conserved sequences (-3, +5) are changed (Reddy, 2007). *In vitro* splicing assays using these pre-mRNA substrates revealed that mutations of the conserved splice site sequences modulate the *in vitro* production of *LHCB3* spliced mRNA (Figure 4B). In agreement with a previous study [66], M1 did not completely abolish the production of spliced mRNA; however, this mutation clearly reduced the generation of the spliced mRNA. Interestingly, M2 completely eliminated the production of the expected product, and instead resulted in a new RNA band of ~100 nt (Figure 4B). These findings further demonstrate that *Arabidopsis* NE supports pre-mRNA splicing.

**Figure 4:**
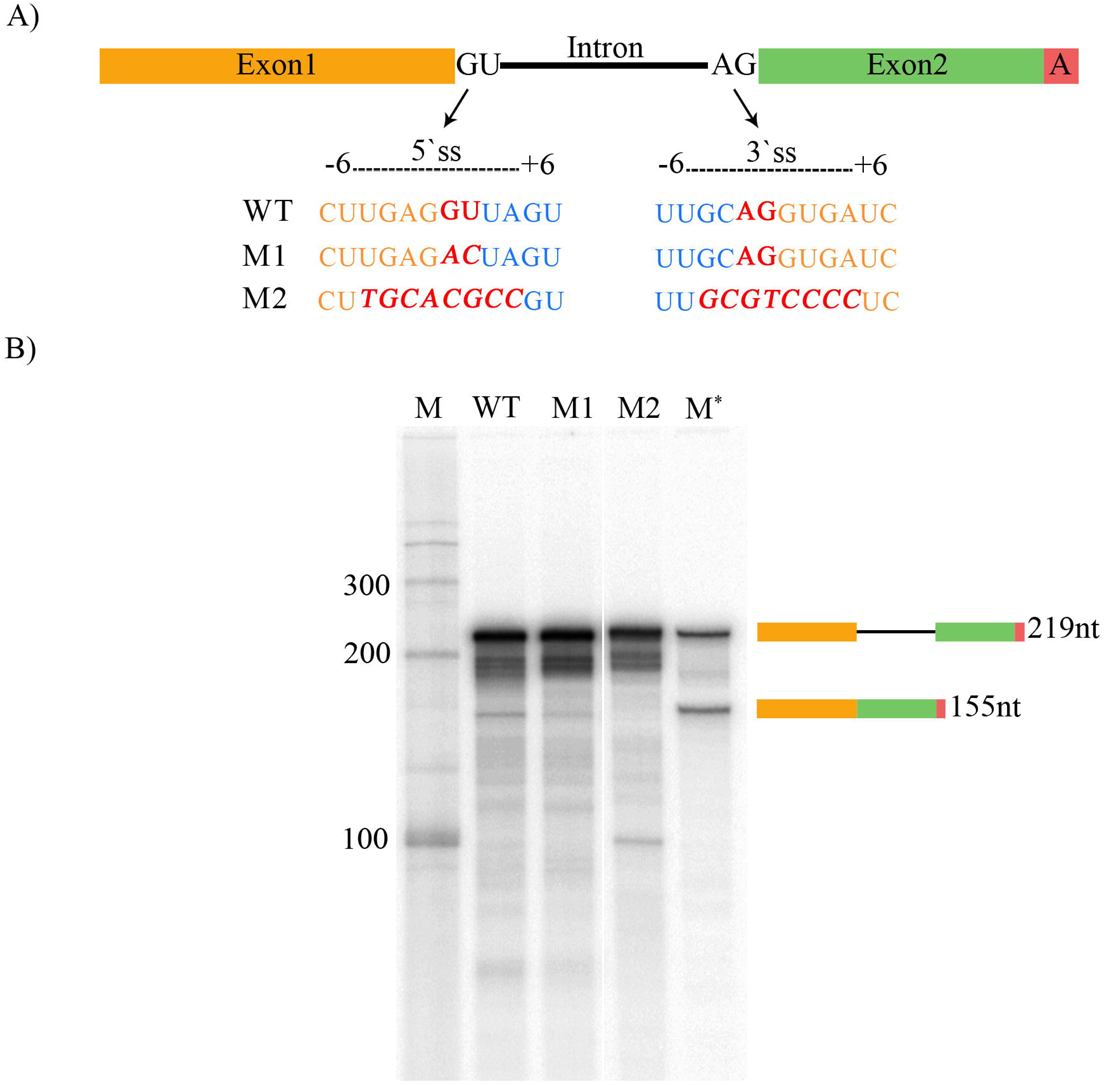
Mutations in conserved splice sites of *LHCB3* pre-mRNA substrate modulated the production of spliced product. (A) Diagram shows sequence substitutions of conserved 5' GU and 3' AG splice sites (ss) of *LHCB3* pre-mRNA substrate. DNA templates with different splice site mutations (M1 and M2) were synthesized (Integrated DNA Technologies, Inc., Coralville, IA) for *in vitro* [^32^P]-labelled RNA synthesis. (B) *In vitro* splicing of [^32^P]-labeled wild type and two mutants (M1 and M2) of *LHCB3* pre-mRNA substrate was carried as described before. Reactions were stopped after three hours. [^32^P]-RNA was isolated and analyzed by electrophoresis as above. RNA markers (M and M*) and schematic diagrams on the right were described in Fig. 2 legend.

### Incubation temperature of *in vitro* splicing reaction affects spliced product

It has been shown previously that different *in vitro* splicing reactions have narrow optimum temperatures. The optimum temperature for the mammalian *in vitro* splicing assay is 30°C, while for the yeast *in vitro* splicing system it is 25°C [2, 4]. Therefore, to address the effect of incubation temperature on the accumulation of the spliced product, reactions were carried out at different temperatures: 24°C, 30°C, 37°C, and 42°C (Figure 5A). The results showed that 24°C is the optimal incubation temperature for high accumulation of the spliced mRNA. The quantity of the spliced product is decreased with increasing incubation temperature. In fact, the incubation at 42°C greatly reduced the appearance of spliced mRNA, suggesting sensitivity of the reaction to high temperature, as was shown in the previous experiment (Figure 2B). Thus, these findings indicate that the *in vitro* splicing assay using *Arabidopsis* NE has an optimal temperature of 24°C, and further demonstrate that the NE contains heat-labile components required for generating the spliced product.

**Figure 5:**
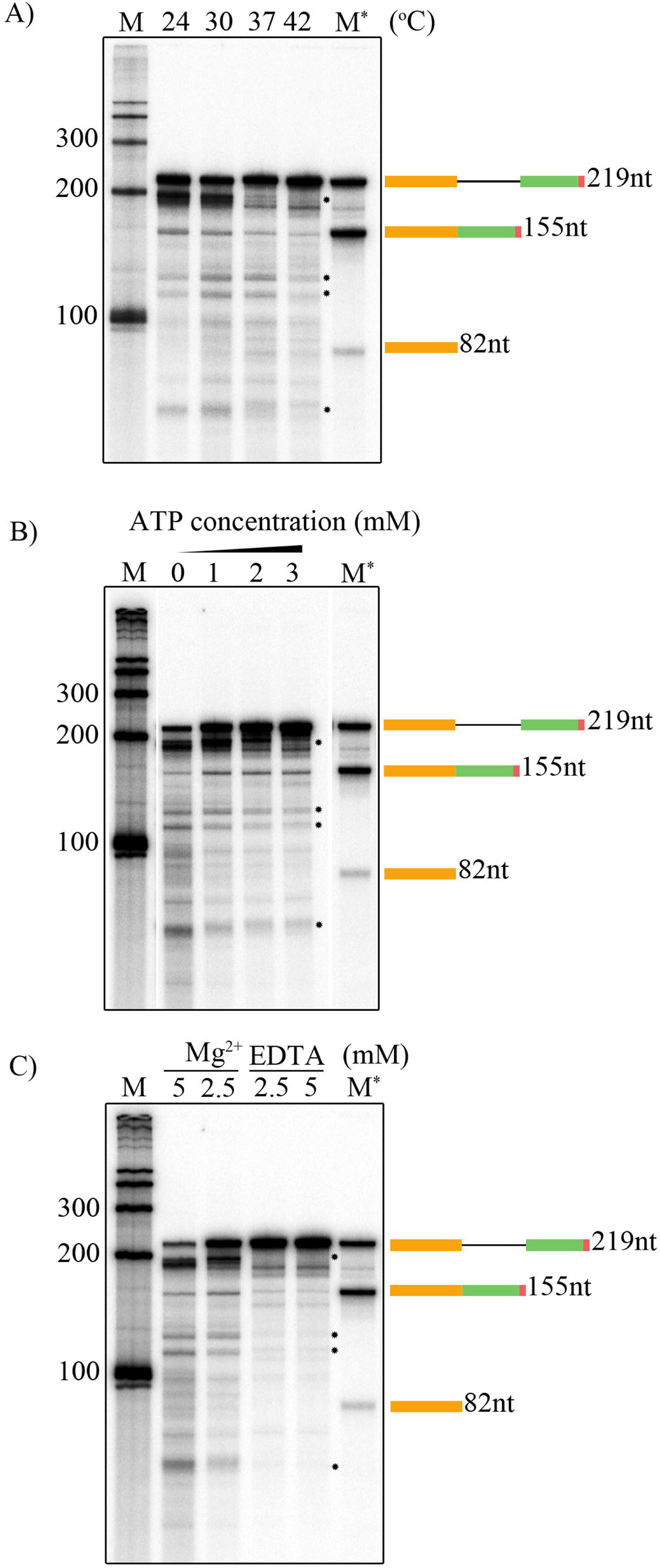
Analysis of optimum conditions for splicing assay. (A) The amount of spliced product at different temperatures. *In vitro* splicing of *LHCB3* [^32^P]-pre-mRNA substrate (25,000cpm) was carried out as described earlier at different temperatures (24°C, 30°C, 37°C, 40°C). (B) Addition of ATP to *in vitro* splicing assay increased the amount of spliced product from *LHCB3* pre-mRNA. (C) Effect of various concentrations of Mg^2+^ on the production of spliced product. *In vitro* splicing reaction of *LHCB3* [^32^P]-pre-mRNA substrate (25,000cpm) was performed as described previously with different concentrations of Mg^2+^ (2.5 and 5 mM), or in the presence of different concentration (2,5 and 5 mM) of EDTA, a divalent cation chelator (EDTA). *In vitro* splicing reaction of *LHCB3* [^32^P]-pre-mRNA substrate (25,000cpm) was carried out as described above without (0mM) with increasing concentrations ATP (1, 2, 3mM). All reactions were stopped after three hours. [^32^P]-RNA was recovered and analyzed by electrophoresis on a 6% polyacrylamide-7 M urea gel, followed by autoradiography. RNA markers (M and M*) and schematic diagrams on the right were described in Fig. 2. The asterisks indicate the potential splicing intermediates. Other [^32^P]-RNA products could be another pre-mRNA splicing intermediates and/or degradation products.

### Effects of ATP and Mg^2+^ on pre-mRNA splicing *in vitro*

It has been well-established that mammalian, yeast, and *Drosophila in vitro* splicing reactions require exogenous ATP and Mg^2+^ [2, 4, 5]. Therefore, we aimed here to investigate and optimize the requirement of these cofactors for the Arabidopsis-derived *in vitro* splicing reaction. Notably, the NE preparation method does not involve a dialysis step against the swelling buffer, hence the NE contains endogenous concentrations of ATP and Mg^2+^. In addition, the 25 μl *in vitro* splicing reaction containing 50% NE has 2.5 mM Mg^2+^, as the swelling buffer contains 5 mM MgCl_2_. The ATP concentration of the *in vitro* splicing reaction was varied while holding the concentrations of remaining components constant (Figure 5B). In the absence of exogenously added ATP, some RNA degradation was observed. In contrast, the addition of ATP maintained input pre-mRNA integrity and also enhanced production of the spliced mRNA. There were no obvious differences in effect between the ATP concentrations (1, 2, and 3 mM) tested. Therefore, these results indicate that the addition of exogenous ATP might support splicing *in vitro,* and the optimal concentration is ~1 mM.

In the same manner, the effect of varying Mg^2+^ concentration on the *in vitro* reaction was investigated (Figure 5C). We found that 2.5 mM Mg^2+^ was likely an optimal concentration for the *in vitro* splicing reaction. Indeed, increasing Mg^2+^ to 5 mM caused input pre-mRNA instability. On the other hand, *in vitro* splicing activity was reduced by the addition of 2.5 or 5 mM EDTA, an ion-chelating agent, indicating a divalent cation requirement (Figure 5C). Taken together, these findings suggest that the plant-derived *in vitro* splicing reaction is like other *in vitro* splicing system requires ATP and Mg^2+^.

## Discussion

*In vitro* splicing systems derived from mammals, yeast, and *Drosophila* have allowed remarkable progress in illustrating splicing mechanisms in eukaryotes. There is no *in vitro* splicing assay for plant systems. Hence, many aspects of pre-mRNA splicing in plants are unknown. In an effort to develop a plant *in vitro* pre-mRNA splicing system, we report here experimental evidence suggesting that NE derived from *Arabidopsis* etiolated seedlings is capable of splicing plant pre-mRNA *in vitro*.

In this study, we show that *Arabidopsis* NE was able to convert the pre-mRNA of *LHCB3* substrate into an expected size of spliced mRNA. The data presented here strongly suggest that the RNA product that corresponds to the spliced mRNA marker is likely a spliced product. Production of the expected size of mRNA upon incubation of pre-mRNA template in the NE, indication that the two exons are linked together according to a junction mapping assay using S1 nuclease, and demonstration that substitution mutations of conserved splice site sequences inhibit the appearance of spliced mRNA – all suggest splicing of *LHCB* pre-mRNA in plant NE. In addition, we found this system to be similar to the well-established non-plant *in vitro* pre-mRNA splicing assays in several ways. First, this system is sensitive to high temperature; second, it requires Mg^2+^; and third, ATP is necessary for the generation of spliced mRNA. From these experiments, we conclude that this is a promising progress towards developing an efficient plant *in vitro* splicing system.

The establishment of an *in vitro* splicing system for plants has been long overdue; difficulties in developing one may be because of characteristics inherent to plant cells. Our success in establishing this *in vitro* assay may be attributed to plant materials used for NE preparation, NE preparation method, and choice of pre-mRNA substrate. The NE is prepared from *Arabidopsis* seedlings that were grown under dark conditions, which are actively growing. The actively growing seedlings have high levels of gene expression for which active pre-mRNA splicing machinery is also needed. Furthermore, unlike other preparation methods that include a high salt lysis buffer, we applied a method that uses low denaturing conditions. This may maintain protein integrity and result in protection of the native states of spliceosomal machinery. Moreover, it is known that not every pre-mRNA substrate can be spliced *in vitro*, and each substrate requires optimized *in vitro* splicing conditions [67]. Thus, the choice of pre-mRNA substrate and its primary structure is also a significant factor for successful *in vitro* splicing systems. We chose *LHCB3* pre-mRNA as this gene is highly expressed and is likely to be processed efficiently. In addition to LHCB3, we tested several other plant pre-mRNAs in plant NE, which are not spliced. In animals also, only a few pre-mRNAs such as β-globin, β-tropomyosin, adenovirus, δ-Crystallin, and Simian virus 40 (SV40) pre-mRNA are widely used, suggesting that only a few pre-mRNA are efficiently spliced under *in vitro* conditions [67-69].

Convincing evidence also suggests that the plant *in vitro* splicing assay system is quite similar to assays used for mammals, yeast, and *Drosophila.* Similarities involve assay conditions, including concentrations of ATP, Mg^2+^, and K^+^ [62]. In addition, the time course of a splicing reaction (0 to 180 min) is comparable between this *in vitro* splicing assay and other such systems [2, 4, 5]. Meanwhile, the optimal incubation temperature for this splicing reaction was unlike other splicing systems; it was within the range of the optimal growth temperature for *Arabidopsis* (23-25^°^C) [70]. Another point of interest is that *in vitro* splicing of the *LHCB3* pre-mRNA substrate generated other RNA species in addition to the spliced RNA. Given the sizes of each part of the pre-mRNA substrate and the production of RNA species within the size range of the intermediates suggest these species are likely splicing intermediates [6, 71]. In addition, the other RNA species that exhibited unexpected mobility on the gel might correspond to lariat-containing RNA intermediates generated during splicing [6, 71]. Meanwhile, we cannot rule out the possibility that some of these RNA species might result from degradation of the pre-mRNA substrate. Another point of interest is provided by the observation that substitution mutations of splice site consensus sequences in the *LHCB3* pre-mRNA modulate the *in vitro* splicing reaction. Some early studies using a point mutation strategy in the mammalian *in vitro* splicing assay have investigated the function of splice site consensus sequences [66, 72]. Compared with their results, our finding that mutation of the 5'ss causes reduction of splicing efficiency indicates some similarities between these two systems. Taken together, these findings strongly support that *Arabidopsis* NE supports *in vitro* splicing system, which could be used to uncover plant-specific splicing regulatory mechanisms.

The efficiency of splicing in this *in vitro* system was low. It is worth mentioning that the efficiency of other initial *in vitro* splicing assays was also low; however, researchers were able to improve the efficiency of *in vitro* splicing assays over time [69, 73, 74]. Further investigations will be very helpful in gaining more insights for improving the efficiency of this plant *in vitro* splicing assay. In addition, it is possible that the concentration of some spliceosomal proteins is less than optimum in our NE, leading to low efficiency. In animals, splicing deficient extracts can be made competent by adding one or more splicing factors such as SR proteins [9, 75]. In future, one could purify one or more SR proteins and supplement the NE or use transgenic lines that are overexpressing one or more SR proteins to prepare NE to enhance splicing efficiency.

## Conclusions

In summary, we show that NE derived from *Arabidopsis* etiolated seedlings is capable of splicing *LHCB3* pre-mRNA. This plant-derived *in vitro* splicing assay system opens new avenues to investigate the spliceosome assembly and composition in plants, splicing regulatory mechanisms specific to plants, and thereby enhance the overall understanding of post-transcriptional gene regulatory mechanisms in eukaryotes.

## Abbreviations

NE: nuclear extract
*LHCB3:*: *LIGHT-HARVESTING CHLOROPHYLL B-BINDING PROTEIN 3*.

## Declarations

## Authors’ Contributions

ASNR conceived the project. MA and ASNR designed experiments and analyzed the results. MA performed all the experiments. MA and ASNR wrote the manuscript.

## Acknowledgements

We thank Drs. Jeffrey Wilusz, Tai Montgomery and Asa Ben-Hur for their comments on this work.

## Competing interests

The authors declare that they have no competing interests.

## Availability of data and materials

All the data and the methods used in this study are available in this published article. Any additional information pertinent to this work can be obtained from the corresponding author.

## Consent for publication

The authors give consent to publish this work.

## Ethics approval and consent to participate

Not applicable.

## Funding

MA is supported by a scholarship from the government of the Kingdom of Saudi Arabia.

## Supporting information

**Figure S1:** Schematic diagram showing the steps in NE preparation from four-day-old *Arabidopsis thaliana* etiolated seedlings. See material and methods for details

**Figure S2:** Sequences of DNA template (A), pre-mRNA substrate (B), and predicted mRNA (C). Arabidopsis LHCB3 (AT5G54270) sequence was extracted from The Arabidopsis Information Resource (TAIR). Exonic sequences are shown in upper case letters, while the intron sequence is shown in lower case. SP6 promoter sequence is highlighted in yellow; primers are highlighted in green with either SP6 or adapter sequences: conserved splicing sites (GT – AG) are highlighted in black, and adaptor sequence is highlighted in blue.

**Figure S3**: (A) Sequences of wild type (WT) and mutated (mutant 1 and mutant 2) DNA templates used to prepare pre-mRNA substrates. B) Oligo DNA sequence used for S1 nuclease assay.

